## Supplementary Material for "A feedforward mechanism for human-like contour integration"

**Deep neural network models provide a powerful experimental platform for exploring core mechanisms underlying human visual perception, such as perceptual grouping and contour integration — the process of linking local edge elements to arrive at a unified perceptual representation of a complete contour. Here, we demonstrate that feedforward, nonlinear convolutional neural networks (CNNs) can emulate this aspect of human vision without relying on mechanisms proposed in prior work, such as lateral connections, recurrence, or top-down feedback. We identify two key inductive biases that give rise to human-like contour integration in purely feedforward CNNs: a gradual progression of receptive field sizes with increasing layer depth, and a bias towards relatively straight (gradually curved) contours. While lateral connections, recurrence, and feedback are ubiquitous and important visual processing mechanisms, these results provide a computational existence proof that a feedforward hierarchy is sufficient to implement gestalt "good continuation" mechanisms that detect extended contours in a manner that is consistent with human perception.**

### SUPPLEMENTARY FIGURES

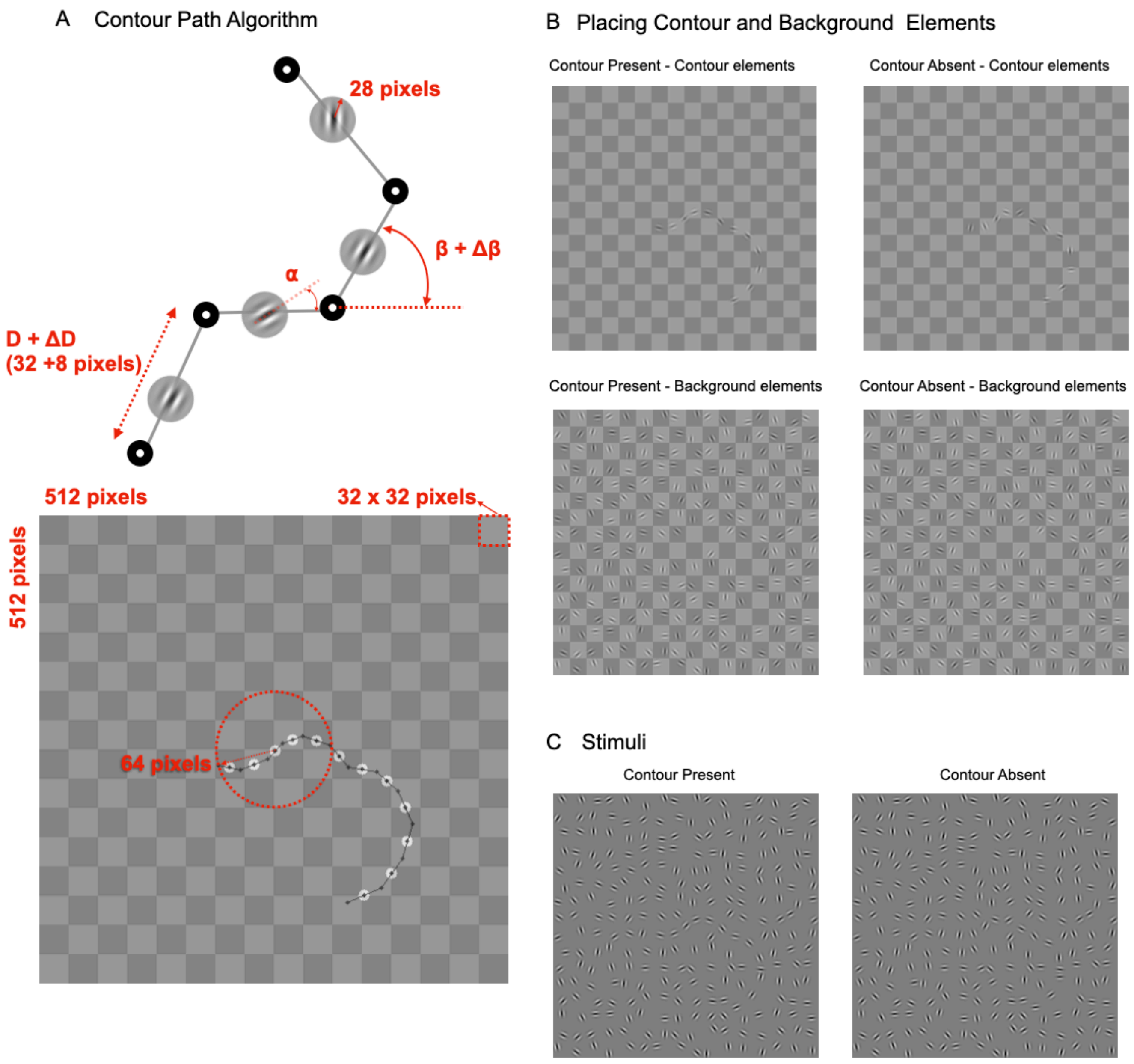


***Supplementary Fig. 1. Contour Stimuli Generation.*** **(A)** Illustration of the contour path algorithm, showing the contour elements (Gabors). These gabors are 28 pixels in size and are about 32 pixels apart with an angular difference of ± (random) β degrees between consecutive contour elements. The bottom panel shows a 512 x 512 pixels grid highlighting the locations of these elements with the first element of the contour always being at a 64 pixel distance from the center. **(B)** Example of placing contour and background elements for contour-present (left) and contour-absent (right) conditions. The top row shows the placement of contour elements and the bottom row shows the placement of background elements. **(C)** Example contour-present and contour-absent stimuli.

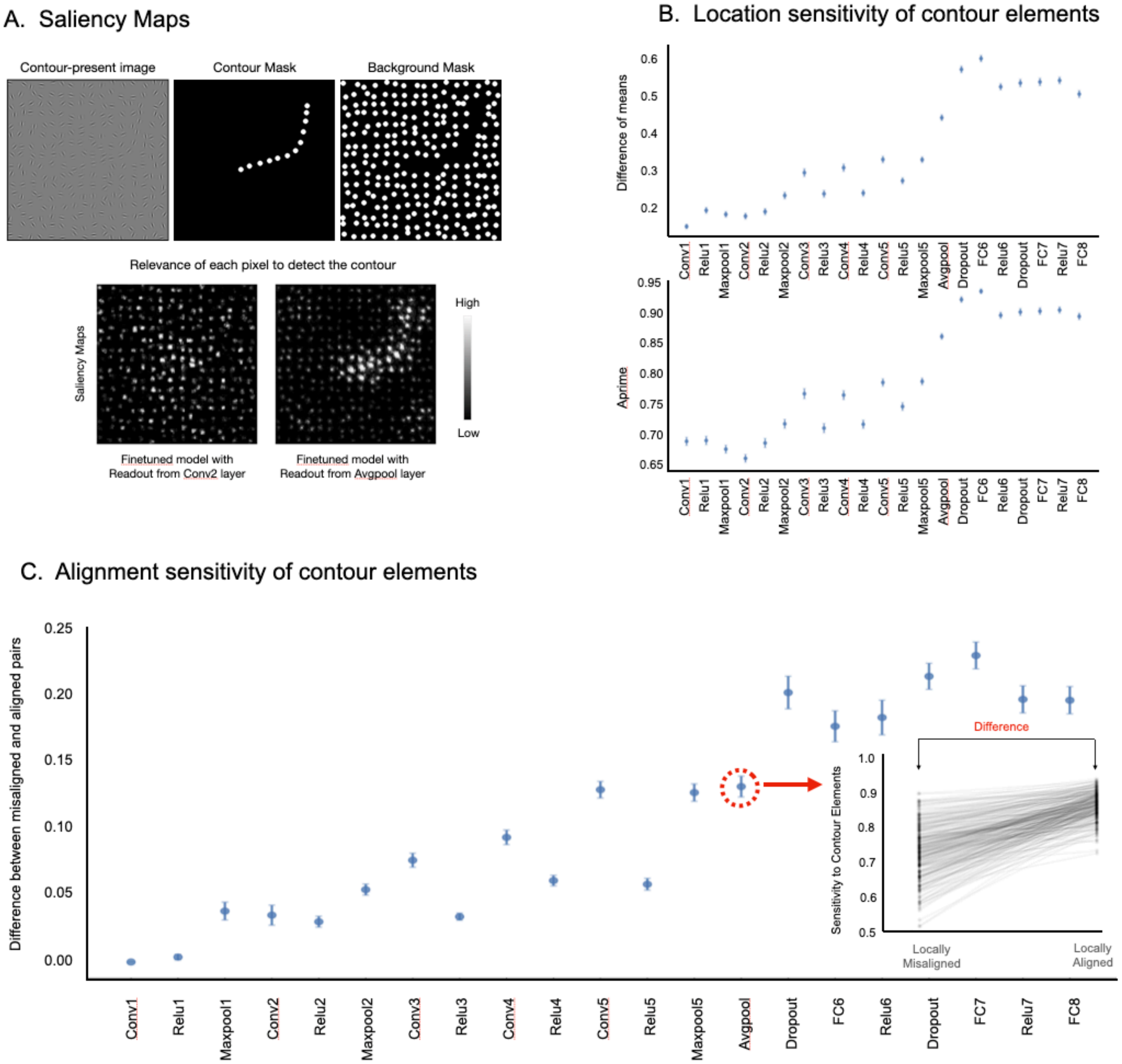


***Supplementary Fig. 2. Sensitivity Analysis of Contour Elements in Fine-Tuned Models***. **(A)** Saliency maps illustrating the relevance of each pixel in detecting a contour within an example image. The top row shows the contour-present image, the corresponding contour mask, and the background mask. The bottom row displays saliency maps for a fine-tuned model with readouts from the Conv2 layer and the Avgpool layer, respectively. **(B)** Location sensitivity of contour elements across different readout layers. The upper plot shows the difference of means, while the lower plot shows aprime (accuracy time) for all readout models. Error bars denote 95% confidence intervals. **(C)** Alignment sensitivity of contour elements, showing the difference between misaligned and aligned pairs across different readout layers. The inset highlights the sensitivity to contour elements (computed using the aprime measure) for locally misaligned and locally aligned elements, with a focus on the difference in sensitivity (indicated by the red arrow and circled region). Error bars represent 95% confidence intervals.

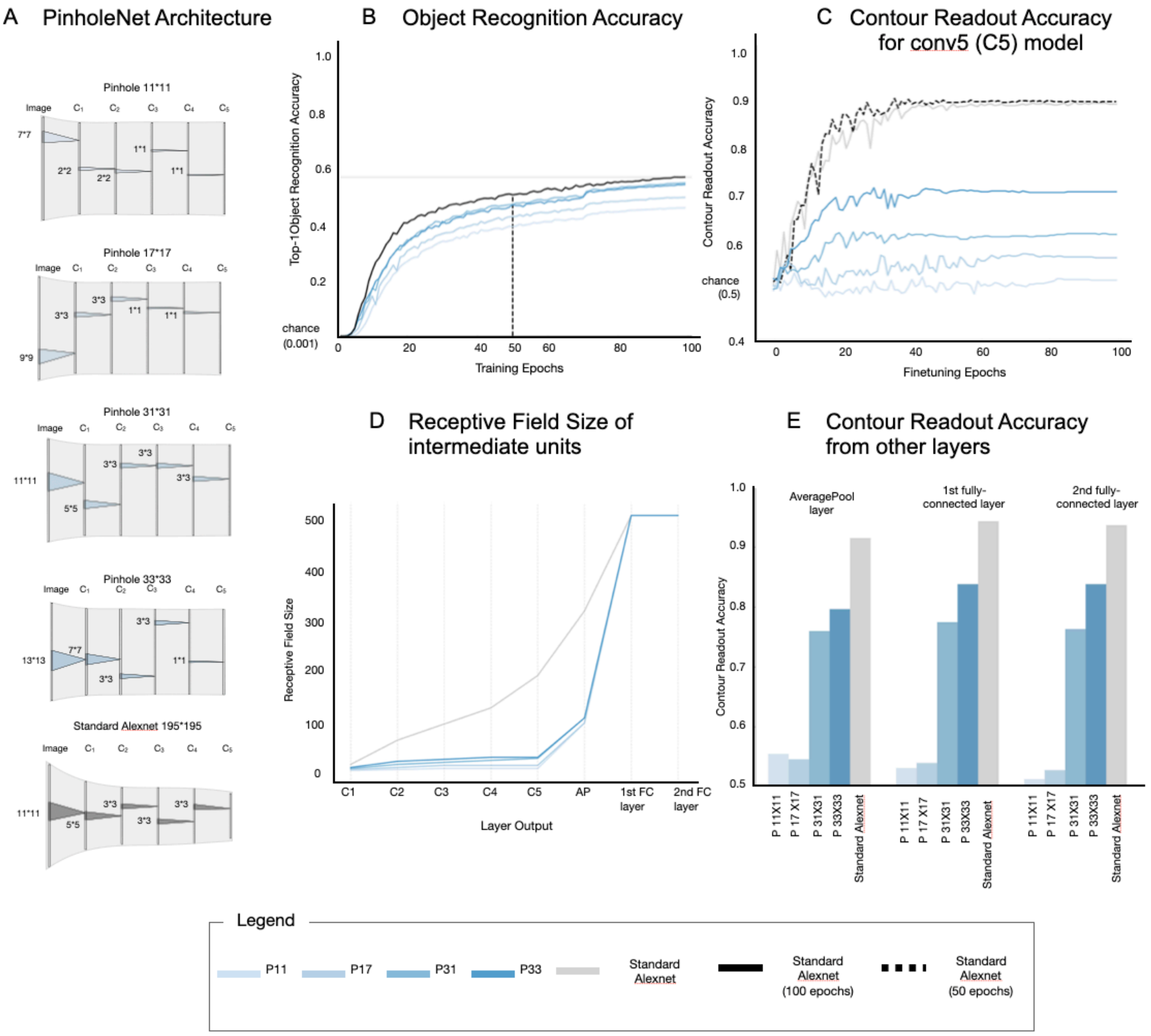


***Supplementary Fig. 3. Architecture and Performance of PinholeNet Architectures.*** **(A)** Schematic representation of PinholeNet architectures with varying receptive field sizes (P11, P17, P31, P33), highlighting the constrained receptive fields in the convolutional layers. For comparison, a standard AlexNet model is also shown in bottom. **(B)** Top-1 object recognition accuracy on the ImageNet validation set during training for different PinholeNet models (P11, P17, P31, P33) and the standard AlexNet model. **(C)** Contour readout accuracy from the Conv5 layer (C5) for models fine-tuned on contour detection. The plot shows the performance of different PinholeNet models compared to the standard AlexNet model across finetuning epochs. **(D)** Receptive field sizes of intermediate units across different layers (C1 to the 2nd FC layer) in PinholeNet and standard AlexNet models. **(E)** Contour readout accuracy from other layers – the avgpool layer, 1st fully connected layer, and 2nd fully connected layer, for different PinholeNet models (P11, P17, P31, P33) and the standard AlexNet model. The legend denotes the color and line style for each model.

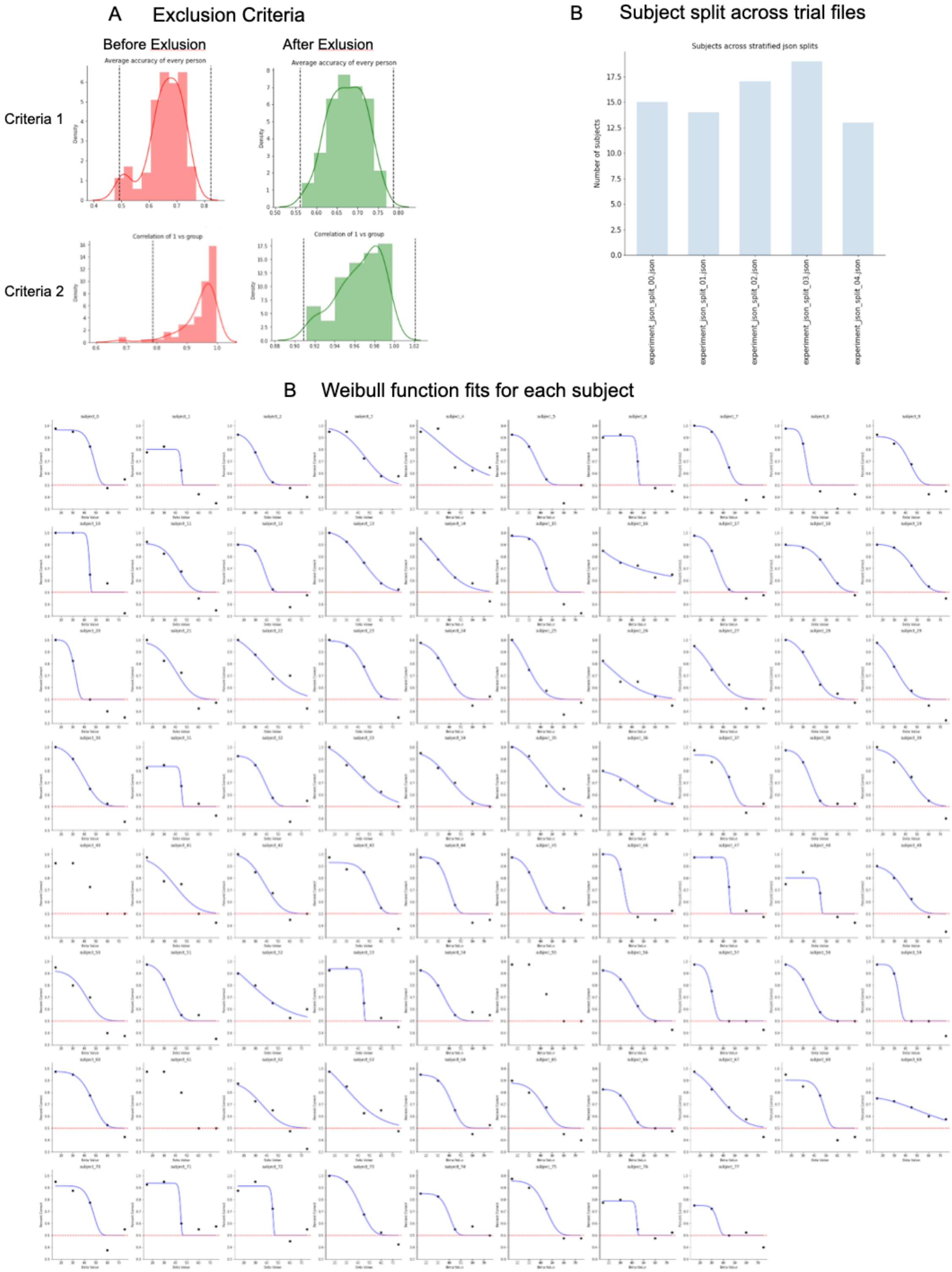


***Supplementary Fig. 10. Human participant performance in Behavioral experiment*** **(A)** Distribution of human performance before and after applying exclusion criteria. The left column shows the performance distribution for Criteria 1 and Criteria 2 before exclusion (red histograms), and the right column shows the performance distribution after applying the exclusion criteria (green histograms). **(B)** Subject split across trial files, indicating the number of participants included in different trial files after applying the exclusion criteria. **(C)** Weibull function fits for each participant, showing individual performance curves. Each subplot represents a different participant, with the x-axis indicating the contour curvature condition and the y-axis representing the performance. The blue line represents the Weibull fit, and the red dashed line indicates chance performance.

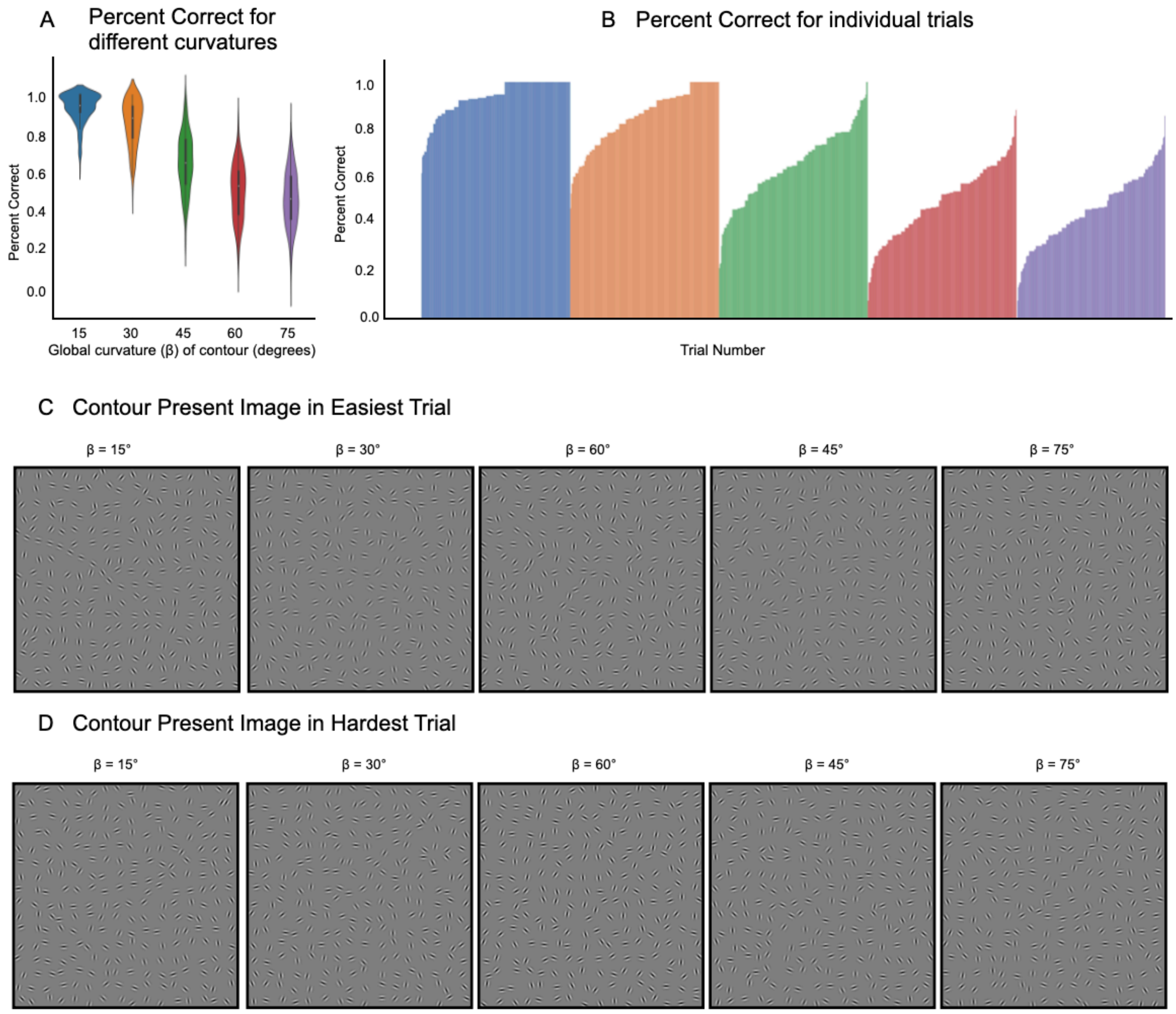


***Supplementary Fig. 5. Human Performance on Contour Detection Across Curvature Conditions.*** **(A)** Violin plots showing the distribution of percent correct responses for different global curvature (β) conditions (15°, 30°, 45°, 60°, 75°). **(B)** Percent correct for individual trials sorted based on global curvature and performance. Each color represents a different global curvature condition. **(C)** Example images showing the contour-present stimulus in the easiest trial for each β condition. **(D)** Example images showing the contour-present stimulus in the hardest trial for each β condition.

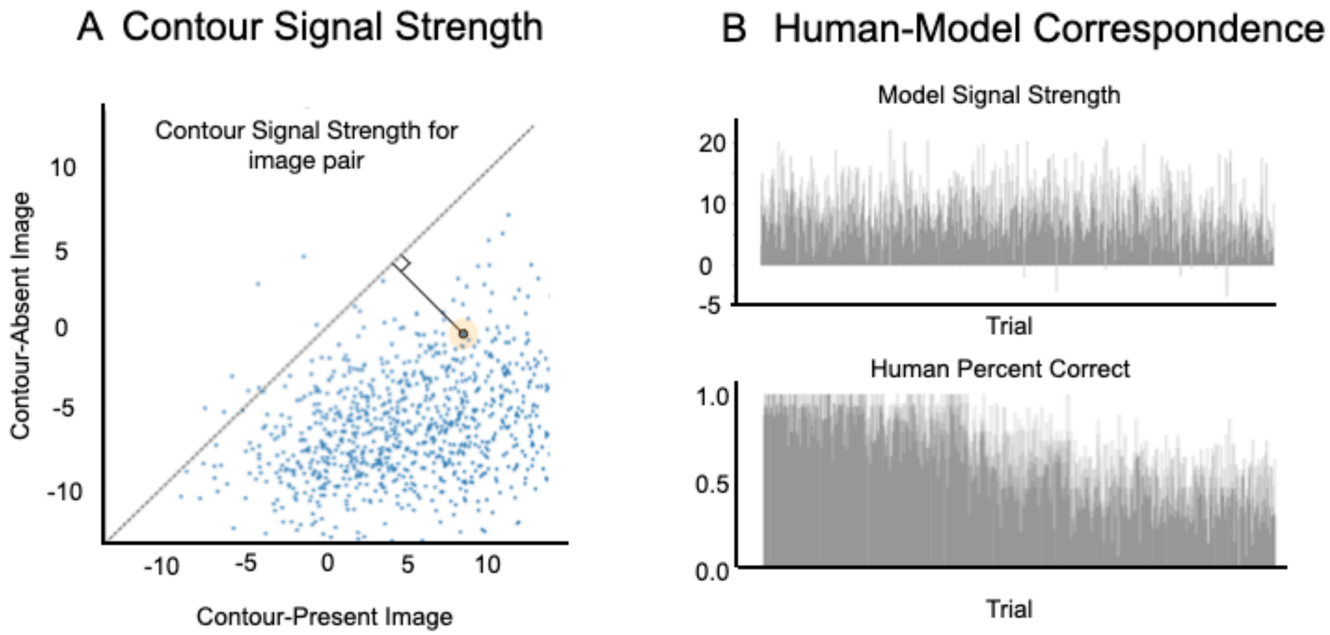


***Supplementary Fig. 6. Analysis of Contour Signal Strength and Human-Model Correspondence.*** (A) Scatter plot showing activity of the contour-present image on the x-axis and the activity of the corresponding contour-absent image on the y-axis. Activity is computed on the contour-present node of the model. Each point represents an image pair of a trial and the perpendicular distance from the diagonal represents contour signal strength for that trial. (B) The top panel shows the model signal strength for each trial, and the bottom panel shows the percent correct responses across humans on the same trials.

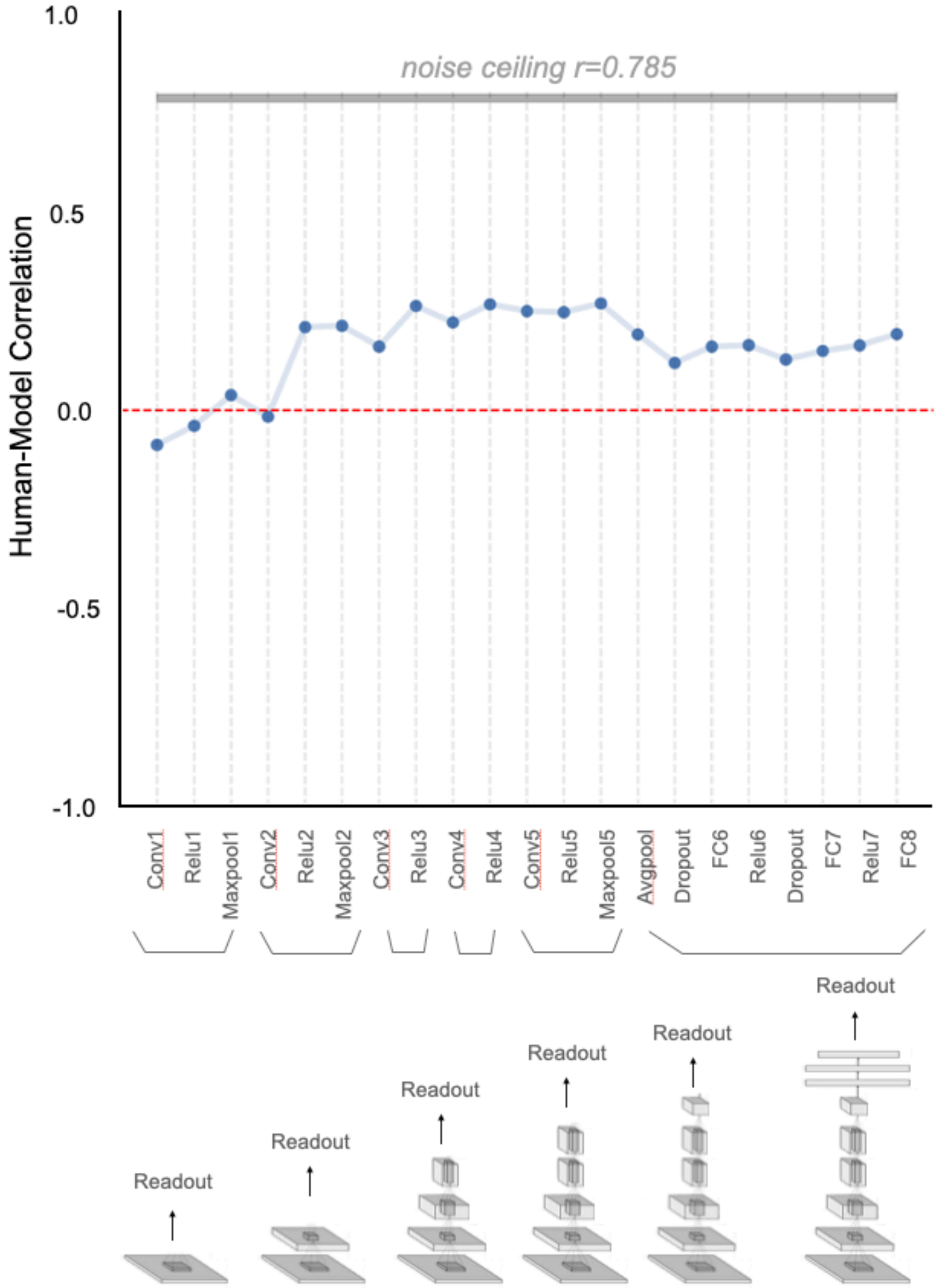


***Supplementary Fig. 7. Human-Model Correlation Across Different Models Reading out from Different Layers of Alexnet and Finetuned on a Broad Range of Contour Curvatures***. The y-axis represents the correlation between human and model contour signal at the level of individual trials, with the noise ceiling indicated by the gray shaded area at r=0.785. The red dashed line represents zero correlation. The x-axis represents the layer from which readouts were taken with a bottom schematic showing the architecture of the contour readout models.

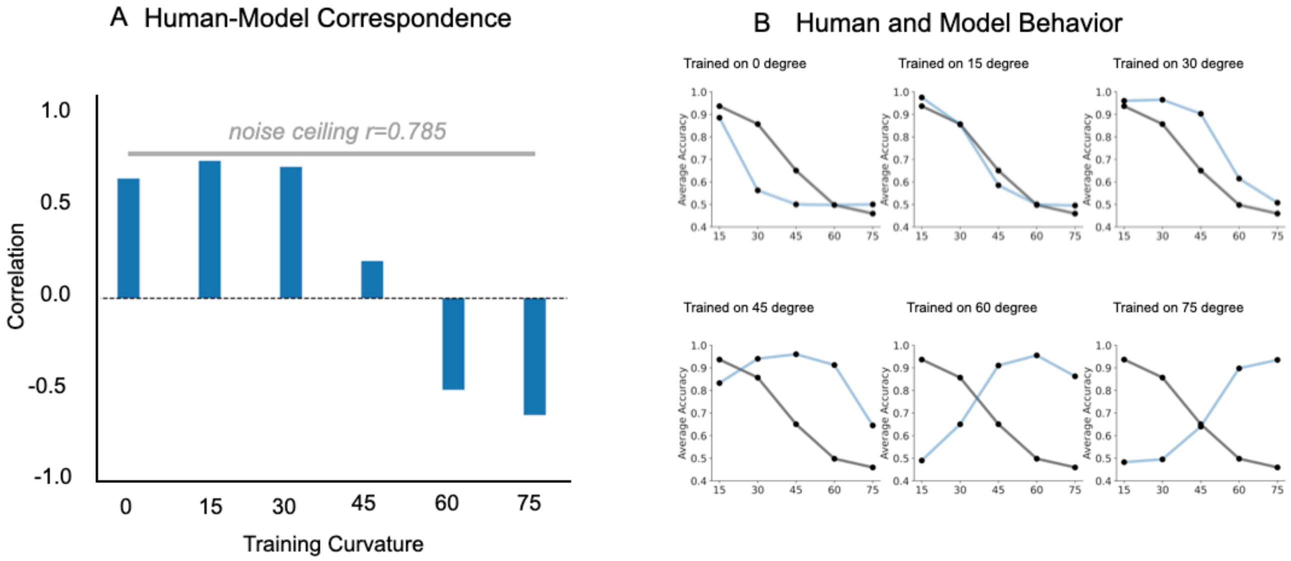


***Supplementary Fig. 8. Human-Model Comparison for Different Readout Models Finetuned on Specific Curvatures (0°, 15°, 30°, 45°, 60°, and 75°).*** **(A)** Human-model correlation of contour signal at the level of individual trials. The y-axis represents the correlation between human and model contour signal at the level of individual trials, with the noise ceiling indicated by the gray shaded area at r=0.785. **(B)** Comparison of humans and model behavior for different global curvature levels. Each subplot shows the performance of the readout model (blue line) and human participants (black line) across varying β conditions.

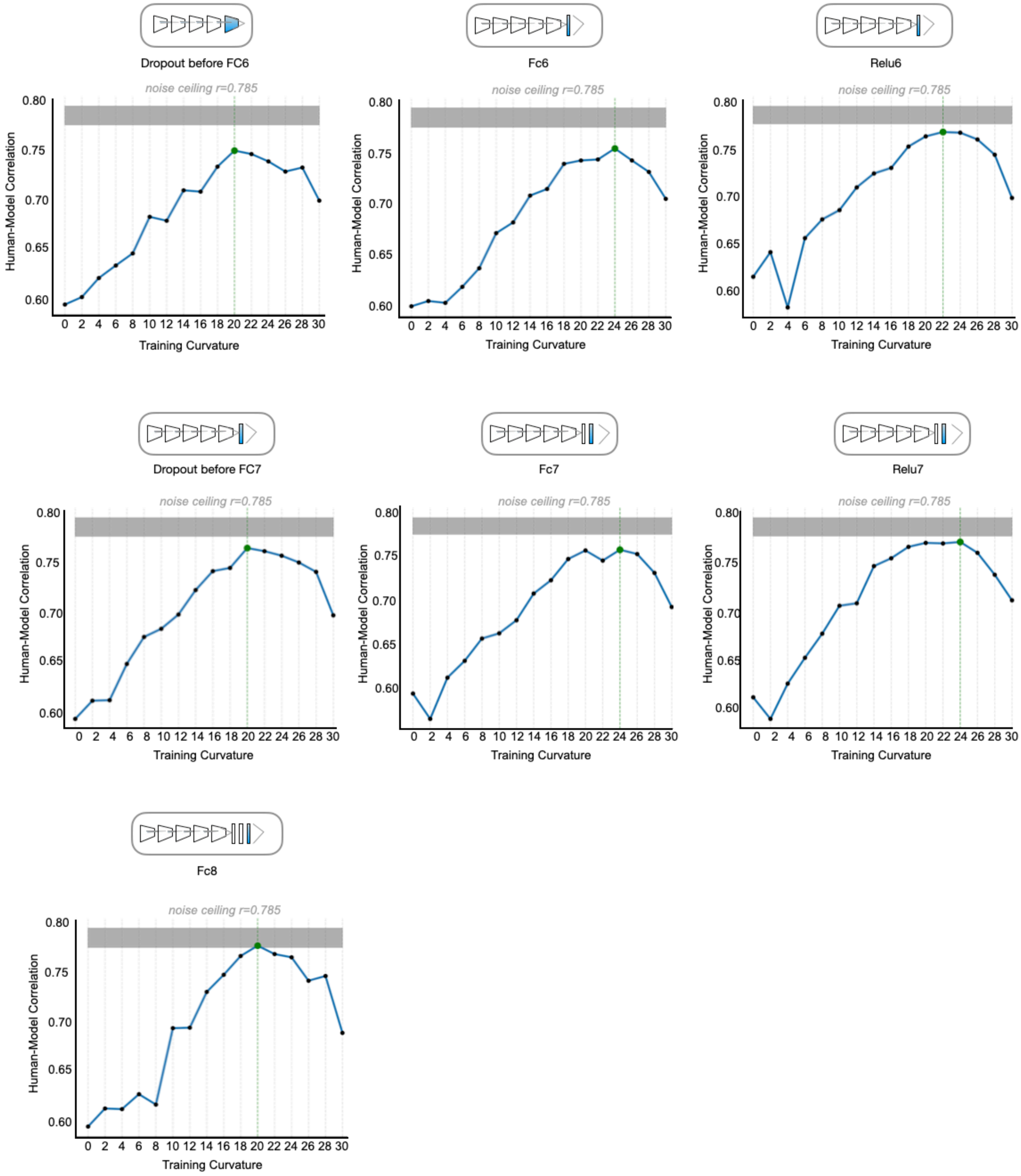


***Supplementary Fig. 9. Human-Model Correlation Across Models Reading out from Different Layers of Alexnet and Finetuned on Specific Contour Curvatures***. Each subplot represents models reading out from a different layer, including Dropout before FC6, FC6, ReLU6, Dropout before FC7, FC7, ReLU7, and FC8. The y-axis represents the correlation between human and model contour signal at the level of individual trials, with the noise ceiling indicated by the gray shaded area at r=0.785. The x-axis represents the training curvature. The schematic above each subplot indicates the architecture of the contour readout models.

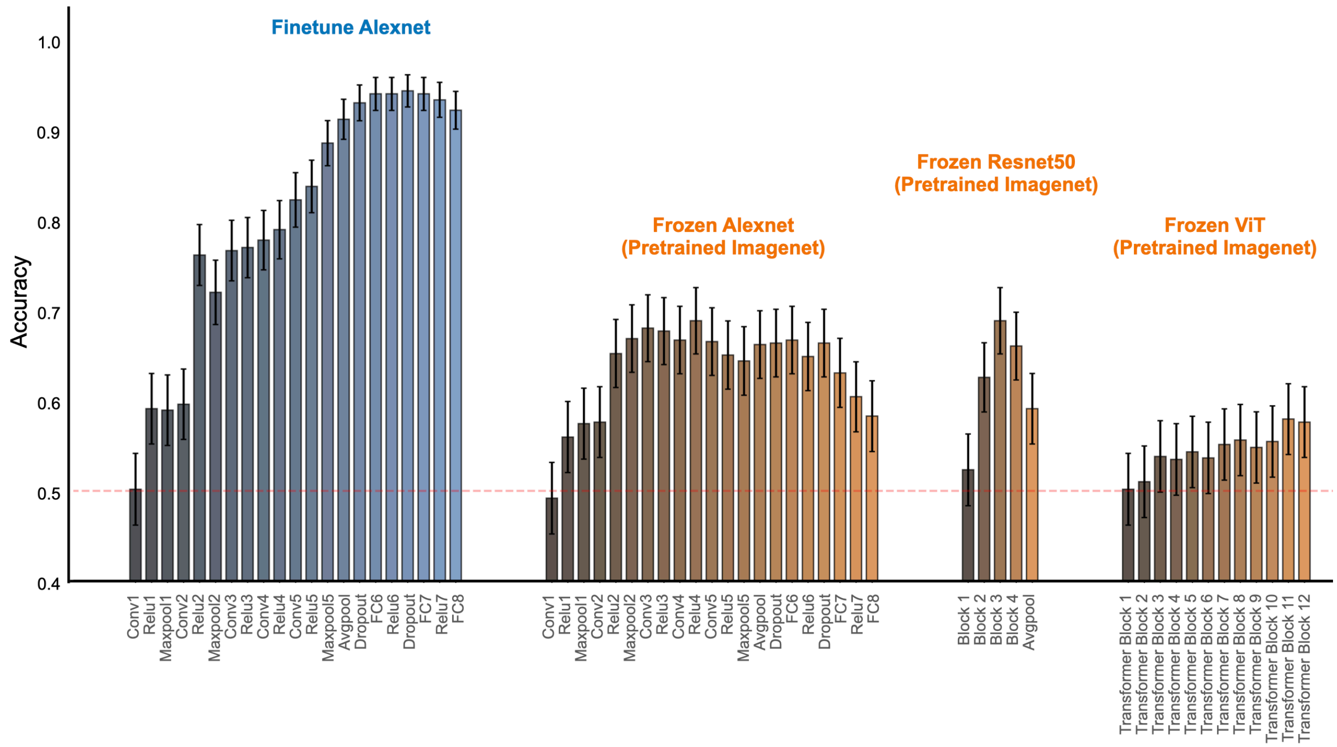


***Supplementary Fig. 10. Contour detection accuracy across different layers of Finetuned Alexnet, Pretrained AlexNet, Pretrained ResNet-50, and Pretrained Vision Transformer models.***. Accuracy values represent the performance of linear readouts trained to detect the presence of contours from the features of each layer. Fine-tuned Alexnet (left, blue) shows significantly improved accuracy across layers, reaching near-perfect performance in the later layers. Frozen AlexNet (pretrained on ImageNet, second from left) shows moderate contour detection performance, peaking in the deeper layers but not matching the fine-tuned variant. Frozen ResNet-50 (pretrained on ImageNet, second from right) and Frozen Vision Transformer (pretrained on ImageNet, right) exhibits similar accuracy trends to the frozen AlexNet model indicating no enhanced contour detection capacity emergent in pretrained networks. Error bars denote 95% confidence intervals for readout accuracy.
